# Lignin accumulation in phloem and outer bark is not associated with resistance to mountain pine beetle in high elevation pines

**DOI:** 10.1101/2021.04.07.438800

**Authors:** David N. Soderberg, Bethany Kyre, Pierluigi Bonello, Barbara J. Bentz

## Abstract

A key component in understanding plant-insect interactions is the nature of host defenses. Research on defense traits among *Pinus* species has focused on specialized metabolites and axial resin ducts, but the role of lignin in defense within diverse systems is unclear. We investigated lignin levels in the outer bark and phloem of *P. longaeva*, *P. balfouriana*, and *P. flexilis*; high elevation species in the western United States known to differ in susceptibility to mountain pine beetle (*Dendroctonus ponderosae*; MPB). Relative to *P. flexilis*, *P. longaeva* and *P. balfouriana* are attacked by MPB less frequently, and MPB brood production in *P. longaeva* is limited. Because greater lignification of feeding tissues has been shown to provide defense against bark beetles in related genera, such as *Picea*, we hypothesized that *P. longaeva* and *P. balfouriana* would have greater lignin concentrations than *P. flexilis*. Contrary to expectations, we found that the more MPB-susceptible *P. flexilis* had greater phloem lignin levels than the less susceptible *P. longaeva* and *P. balfouriana*. No differences in outer bark lignin levels among the species were found. We conclude that lignification in *Pinus* phloem and outer bark is likely not adaptive as a physical defense against MPB.

## Introduction

Bark beetles (Coleoptera: Curculionidae, Scolytinae) are key forest disturbance agents globally and include many tree-killing species [1]. Overcoming tree defenses is a central challenge for bark beetles which feed on living phloem and requires the destruction of tree vascular tissue for offspring survival. Tree defenses serve to protect against insect infestation, thereby maintaining the functional integrity of two subcortical high-fitness-value tissue types: phloem, which is responsible for transport and distribution of photosynthate produced in leaves and needles; and xylem, which provides structural support and functions in translocation of water and dissolved minerals from roots to the rest of the tree. Both tissue types also play a role in defense against bark beetles [2,3] and their fungal mutualists [4–6].

An ecologically and economically significant bark beetle with an extensive distribution across western North America is the mountain pine beetle (MPB) (*Dendroctonus ponderosae* Hopkins, Coleoptera: Curculionidae, Scolytinae) [7,8]. While the majority of *Pinus* species are considered MPB hosts [9], successful MPB attacks on *P. longaeva* (Great Basin bristlecone pine) and *P. balfouriana* (foxtail pine) are rare [10], relative to the commonly attacked *P. flexilis* (limber pine) [11–13]. In addition, MPB displays aversion to *P. longaeva* in both field [14] and laboratory settings [15], and extremely few MPB offspring emerge from manually-infested *P. longaeva* relative to *P. flexilis* [16]. *P. longaeva* and *P. balfouriana* also have dense sapwood and heartwood and possess high concentrations of constitutive specialized metabolite defense compounds relative to co-occurring *P. flexilis* [10].

Specialized metabolites as well as anatomical structures are fundamental in conifer defense. They can be expressed constitutively or upregulated upon attack as needed to maximize the economy of available resources [17–20]. Variation among and within conifer species in chemical [10,21,22] and anatomical defenses [23,24] is well known and hypothesized to reflect resistance to multiple bark beetle species [25,26]. Specialized metabolites include low molecular weight (LMW) compounds (e.g., terpenes and their derivatives, phenolics) that can be toxic to attacking bark beetle adults [27–29] and their eggs and larvae [30], and inhibit the propagation of fungal symbionts [31]. Anatomical defenses are structural elements (e.g., resin ducts, lignified stone cells) that can deter invading insects by providing physical and chemical barriers to nutrient-rich tissues [20,32,33].

Lignin, a fundamental plant structural element, is the second most naturally abundant biopolymer in plant cell walls, after cellulose [34,35]. Lignin is deposited in the secondary cell wall of all vascular plants [36,37] where it provides rigidity for structural stability and impermeability for more efficient water transport [38], as well as structural resilience against abiotic stressors [39–42]. Lignin also plays a role in tree defense, where it can increase resistance to degradation by microorganisms [43–45], and provide protection against pathogenic fungi [46,47] and bacteria [48]. Cell wall lignification also confers tree resistance against herbivory in the form of indirect chemical defenses (i.e., antifeedant or antinutritional) [49,50] and direct physical defenses [51].

In the family *Pinaceae*, sclerenchyma cells of the phloem occur as large stone cells that are primarily comprised of lignin [32,52,53]. Increased stone cell frequencies within the phloem of Sitka spruce (*Picea sitchensis* Bongard) were associated with decreased spruce weevil (*Pissodes strobi* Peck) growth rate, survival, and fecundity, and disruption of larval establishment [53–56]. Decreased growth rate and survival of great spruce bark beetle (*Dendroctonus micans* Kugelann) larvae were also associated with increased lignin concentrations [32,57] and naturally occurring compounds originating from lignin were found to have antifeedant effects on another bark beetle, *Hylobius abietis* (L.) [58]. Moreover, lignin synthase genes were found to be more prevalent in spruce that were beetle-resistant [53]. Because lignified tissue is difficult to chew and digest [32,53,59] it can reduce nutritional quality and nutrient bioavailability [51,60,61] by preventing adequate feeding and increasing mandibular wear [32].

The genus *Pinus,* specifically, is known to have evolved various defensive strategies against phloem-feeding bark beetles, such as specialized metabolites [62–64] and resin ducts [20,65,66], both of which show high variability within and among *Pinus* species [67,68]. Little is known, however, of the role of lignin as a constitutive defensive mechanism against bark beetle attacks in high elevation pine species in the western United States that are at increasing risk due to climate change. We attempted to fill this gap by quantifying lignin in the outer bark (i.e., rhytidome) and phloem of co-occurring *P. longaeva, P. balfouriana* and *P. flexilis* from multiple sites and compared concentrations within and among species and between the two tissue types. We hypothesized that the more MPB-resistant *P. longaeva* and *P. balfouriana* would have greater lignin concentrations than co-occurring *P. flexilis*.

## Methods

### *Site Selection* and *Tree Sampling*

Between June and September 2016, trees were sampled at five sites across the ranges of *P. longaeva* and *P. balfouriana*, four of them in stands with co-occurring *P. flexilis* (Fig 1; Table 1). Four of the five sites were also sampled by Bentz et al. (2017) [10], allowing a comparison with results from that study. Equal numbers of *P. longaeva* and *P. flexilis* trees were sampled at three geographically separated locations, and equal numbers of *P. balfouriana* and *P. flexilis* were sampled at the Sierra Nevada site. At the Klamath site *P. flexilis* was not present, and only *P. balfouriana* was sampled. At each site 15 live trees of each species were sampled, and diameter at breast height (DBH, ~ 1.5 m above ground) ranged from 30-46 cm. Study sites without signs of MPB or pathogen activity were chosen to avoid an influence of induced defenses.

**Fig 1.**
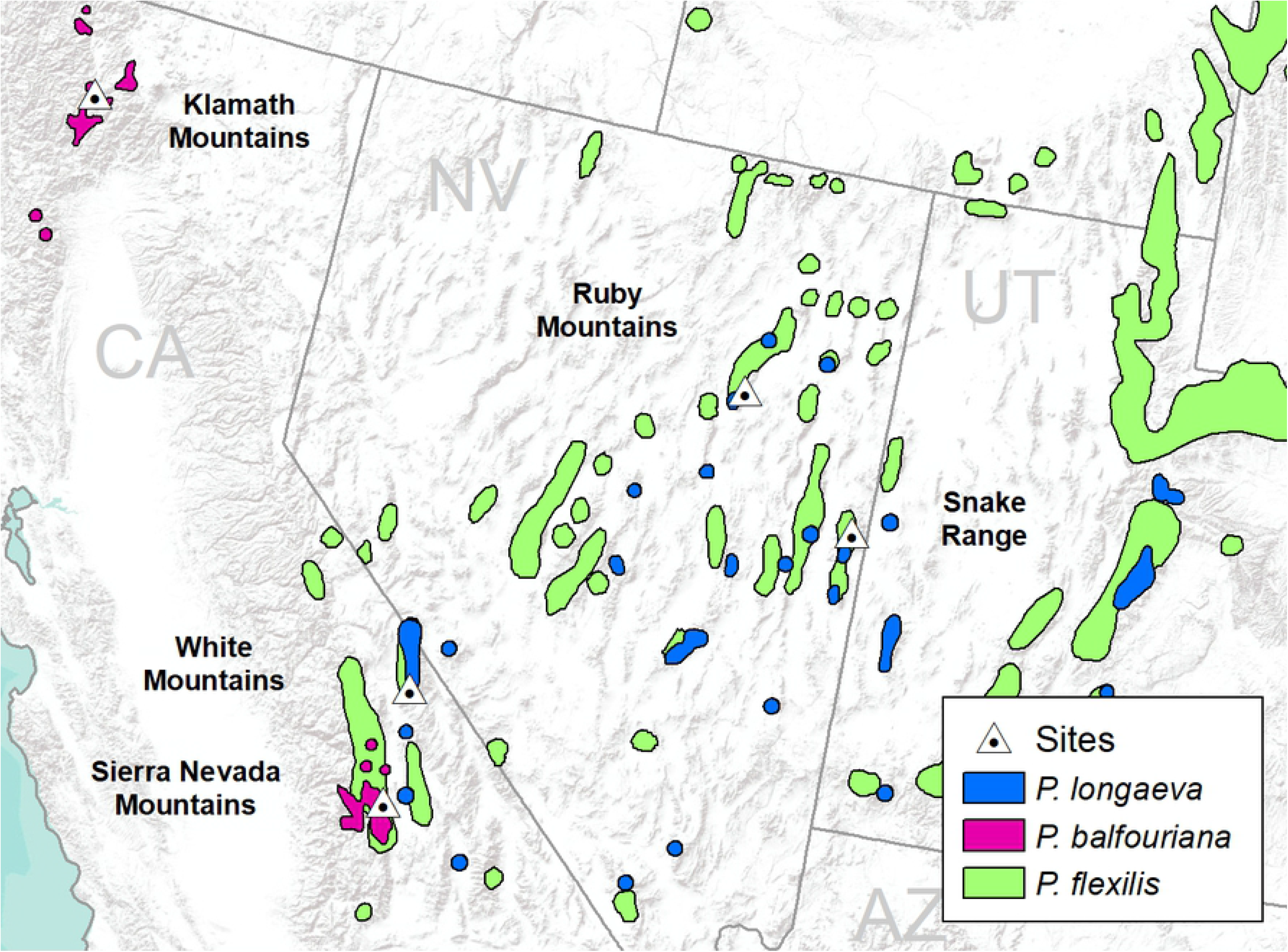
Distributions of Great Basin bristlecone pine (*Pinus longaeva*), foxtail pine (*P. balfouriana*), and limber pine (*P. flexilis*), and sample site locations (see Table 1). Pine distributions are based on Little (1971) [82].

**Table 1.**
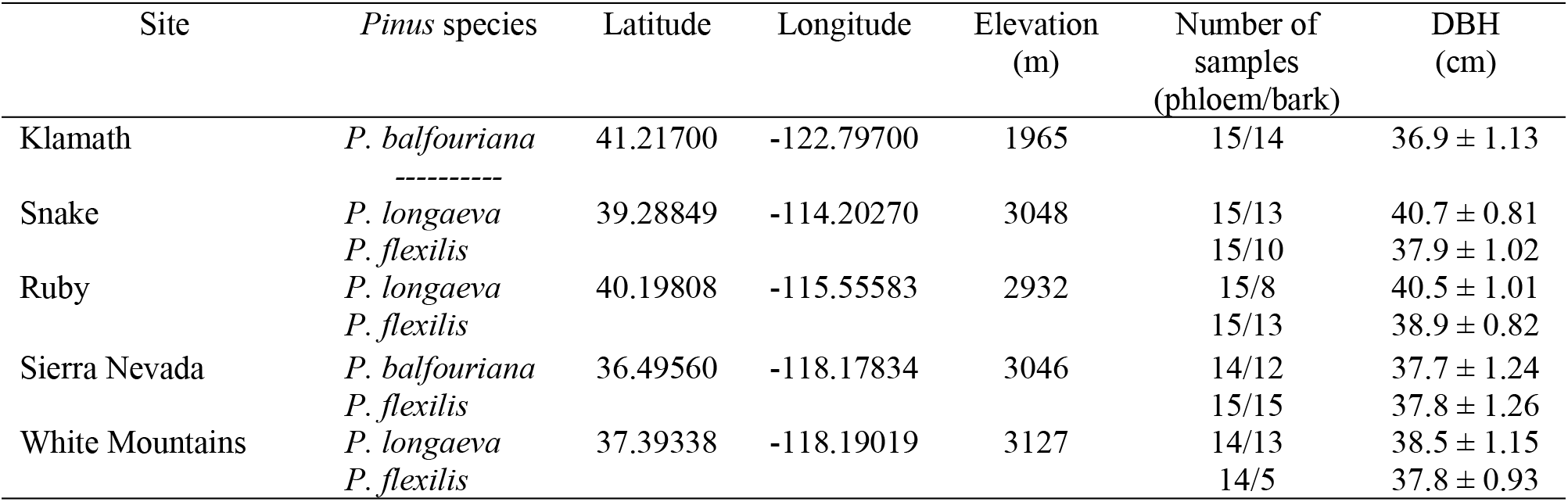
Site locations (see Fig. 1) and stand metrics including species sampled, number of phloem and bark samples analyzed, and mean ± standard error of DBH (diameter breast height).

To assess lignin levels (mg/g FW) in outer bark and phloem, trees were sampled by boring into the tree at breast height with a 1” diameter circular hole saw. Four samples were taken on the north, south, west, and east facing aspects of the tree trunk and pooled to account for potential within-tree variation. Upon tissue removal, phloem thickness (mm) was measured from the north and south aspect samples. Outer bark and phloem tissues were then separated and placed immediately in a sealed vial in a cooler with dry ice for transport to the Rocky Mountain Research Station (Logan, UT) for cold storage (−40**°**C).

### Lignin extraction

In the laboratory, outer bark and phloem samples were prepared for lignin extraction using a ceramic mortar and pestle to grind tissue samples in liquid nitrogen. Tissues were ground to a fine powder and placed in vials for lignin extraction. The mortar and pestle were cleaned with 95% ethanol between each tissue sample. Lignin was extracted from the outer bark and phloem tissues using thioglycolic acid digestion in a modification of the method of Bruce and West (1989) [69], as described by Bonello et al. (1993) [70]. Spectral absorbance of phloem lignin samples (n = 135) was measured at 280 nm using a NanoDrop™ 3300 Fluorospectrometer (Thermofisher Scientific) with a 1:4 dilution in NaOH against a standard curve of pure spruce lignin (Sigma-Aldrich) at 0, 18, 45, 90, and 360 micrograms/mL. The spectral absorbance of outer bark lignin (n = 103) was measured under the same parameters using 1:64 dilution. All phloem samples were assessed as pure and free from contamination, although thirty-two outer bark samples were removed from analysis due to residual phenolic compound contamination (S1 Fig). In addition, three outliers, consisting of a single phloem sample from each species (2% of total samples), exhibited lignin concentration > 6-fold the standard deviation for each species. As the outer bark contained remarkably higher lignin concentrations than the phloem, we removed these three outliers out of caution for potential tissue contamination. Adjusted sample sizes for outer bark and phloem samples are shown in Table 1.

### Statistical Analysis

Differences among tree species in phloem and outer bark lignin concentrations, phloem thickness, and DBH were assessed with a hierarchical mixed effect analysis of variance (ANOVA), that accounts for variation among sites, using the package “lme4” [71] in R version 4.0.0 [72]. Multiple comparisons among sites were assessed using the package “multcomp” [73]. Linear regression (package “lme4”) was used to assess the relationships between phloem and outer bark lignin concentrations, phloem lignin concentration and phloem thickness, DBH and phloem thickness, DBH and phloem lignin concentration, and DBH and outer bark lignin concentration.

## Results

Phloem lignin concentrations did not differ between *P. longaeva* and *P. balfouriana*, but, contrary to our hypotheses, *P. flexilis* had significantly higher (~2-fold) phloem lignin concentrations than the other two species (Fig 2; Table 2). We found no differences among the species in outer bark lignin concentrations (Fig 2; Table 2). *P. flexilis* had thinner phloem than both *P. longaeva* and *P. balfouriana*, but there were no differences in phloem thickness between *P. longaeva* and *P. balfouriana* (Fig 3; Table 2). *P. flexilis* trees with thicker phloem tended to have lower phloem lignin levels, but we found no relationship between phloem thickness and phloem lignin levels in *P. longaeva* or *P. balfouriana* (Table 3). We also found no relationship between phloem thickness and outer bark lignin levels in any species (Table 3)*. P. flexilis* and *P. balfouriana* were generally smaller than *P. longaeva* (Table 2), although DBH had no effect on phloem or lignin concentrations in any of the species (Table 3). There was also no significant relationship between phloem and outer bark lignin concentrations among trees, although *P. balfouriana* with more phloem lignin tended to have less outer bark lignin (Table 3). There were no significant differences among the sites in phloem lignin concentrations for any species (*P. flexilis*: p > 0.238; *P. longaeva*: p > 0.095; *P. balfouriana*: p = 0.101), although *P. flexilis* outer bark lignin concentration differed at two sites (S1 Table).

**Fig 2.**
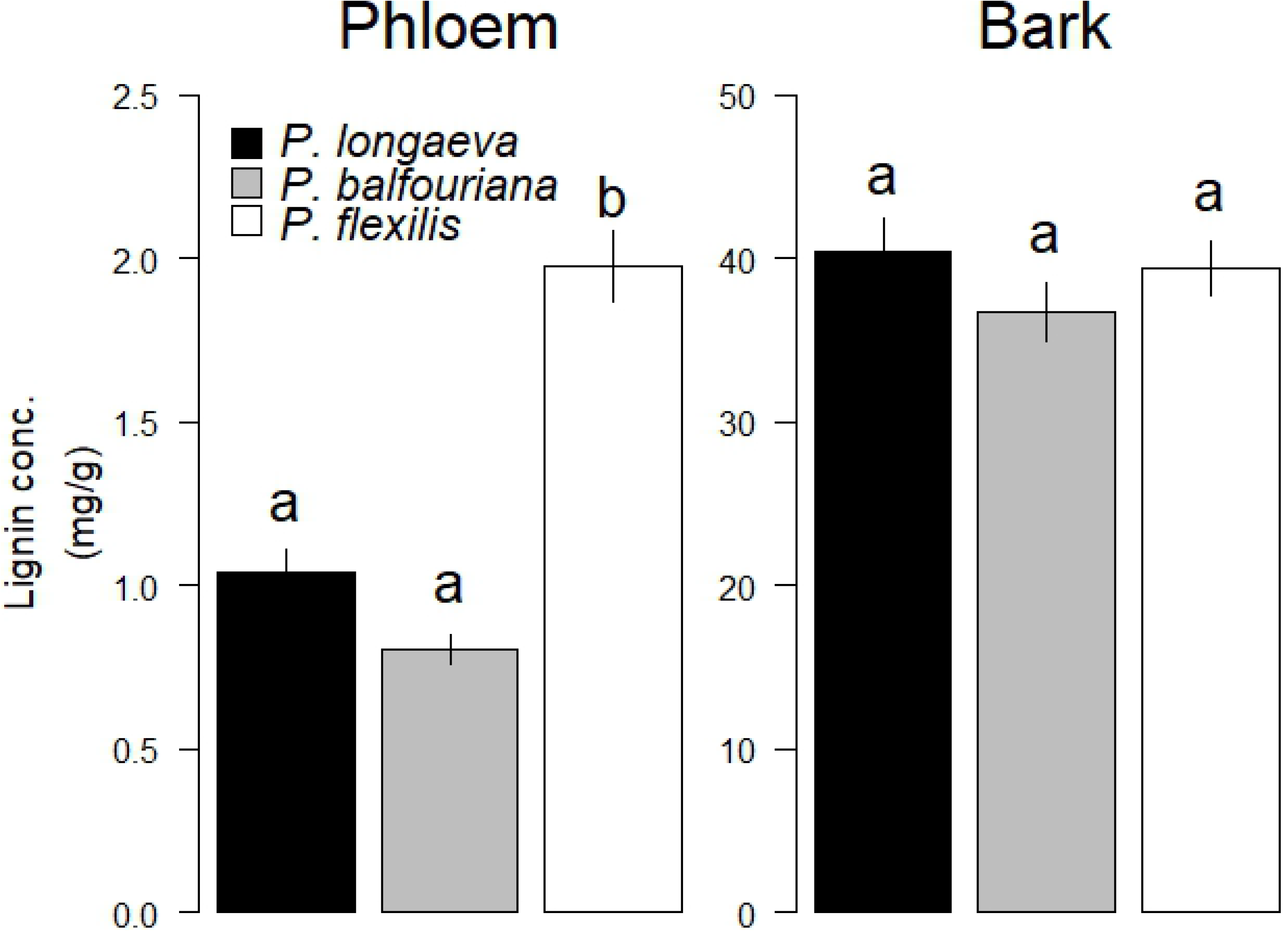
Phloem and bark lignin concentratons (± standard error) in *P. longaeva, P. balfouriana,* and *P. flexilis* averaged across all sites. Different letters (i.e., a,b) denote statistically significant differences among species means (p < 0.05). See Table 2 for statistics.

**Fig 3.**
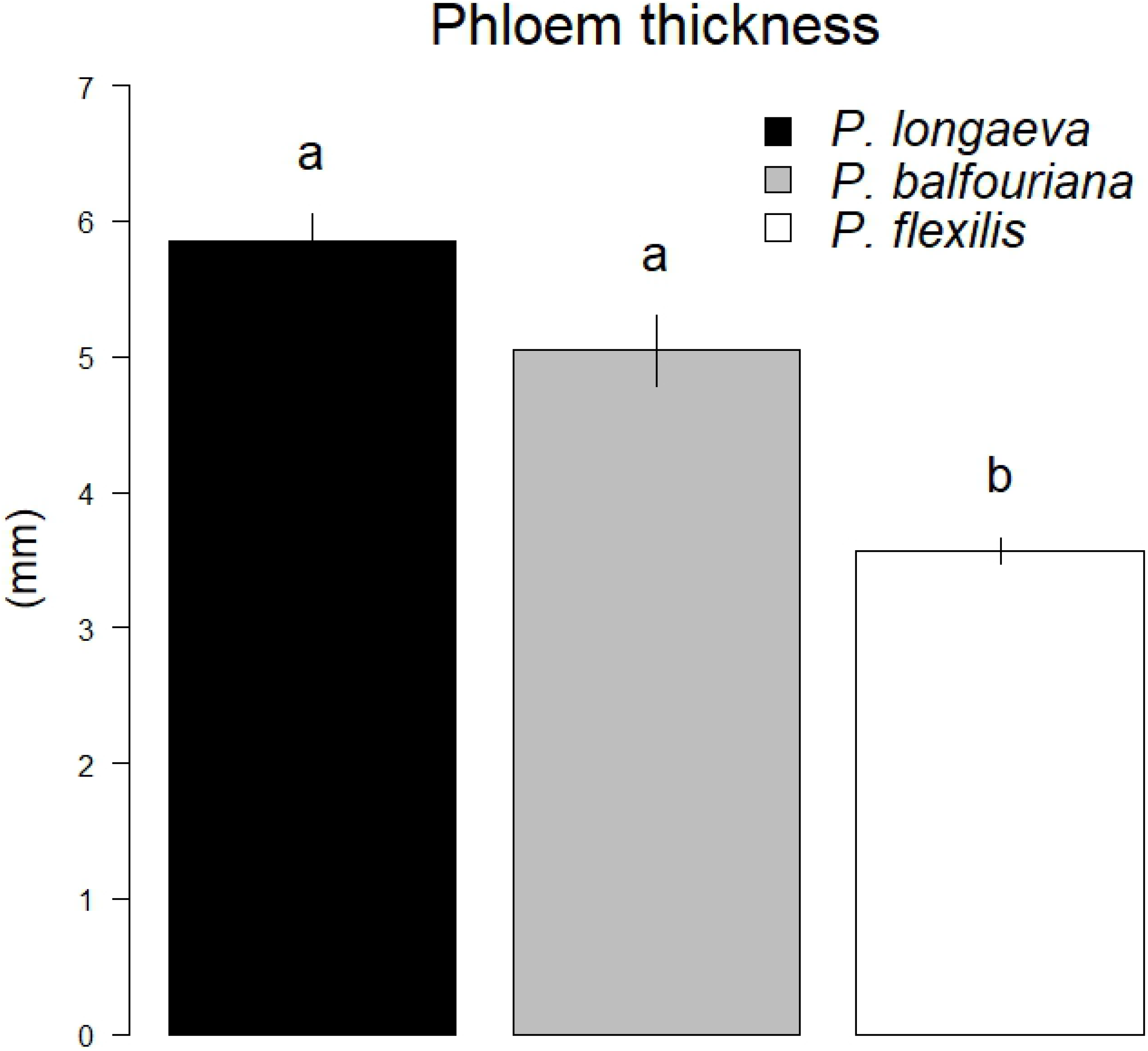
Phloem thickness (± standard error) in *P. longaeva, P. balfouriana,* and *P. flexilis*, averaged across all sites (see Fig. 1; Table 1). Different letters (i.e., a,b) denote statistically significant differences among species means (p < 0.05). See Table 2 for statistics.

**Table 2.** Model estimates testing for species differences in diameter at breast height (DBH; cm), phloem thickness (mm), and phloem and bark lignin concentrations (g/mg FW) among *P. flexilis*, *P. balfouriana*, and *P. longaeva*. Effect size (Est.) and 95% confidence interval (95%CI) estimates between comparison samples are shown. P-values (p) describe the likelihood of statistical difference with values < 0.05 presented in bold.

|  | DBH |  | Phloem thickness |  | Phloem lignin |  | Bark lignin |  |
| --- | --- | --- | --- | --- | --- | --- | --- | --- |
|  | Est. (95%CI) | p | Est. (95%CI) | p | Est. (95%CI) | p | Est. (95%CI) | p |
| *P. flexilis* vs *P. balfouriana* | 0.71 (-1.76, 3.19) | 0.780 | -1.83 (-2.73, -0.03) | **<0.0001** | 0.91 (0.47, 1.33) | **<0.0001** | 3.04 (-4.91, 11.0) | 0.641 |
| *P. flexilis* vs *P. longaeva* | -2.19 (-4.40, 0.01) | 0.052 | -2.27 (-2.89, -1.65) | **<0.0001** | 1.03 (0.75, 1.32) | **<0.0001** | -1.54 (-7.58, 4.51) | 0.821 |
| *P. balfouriana* vs *P. longaeva* | -2.91 (-5.38, -0.43) | **0.016** | -0.44 (-1.48, 0.61) | 0.584 | 0.13 (-0.36, 0.62) | 0.744 | -4.57 (-13.5, 4.32) | 0.448 |

**Table 3.** Modeled linear regression coefficients (i.e., slope) testing for the relationship between phloem thickness (mm) and DBH (diameter at breast height, cm), phloem thickness and phloem lignin concentrations (g/mg FW), phloem lignin concentrations and DBH, bark lignin concentrations and DBH, and phloem and bark thickness within *P. flexilis*, *P. balfouriana*, and *P. longaeva* across sites. Effect size (Est.) and 95% confidence interval (95% CI) estimates between comparison samples are shown. P-values (p) presented describe the likelihood of statistical difference with values < 0.05 presented in bold.

|  | Phloem thickness × DBH |  | Phloem thickness × Phloem lignin conc. |  | Phloem lignin conc. × DBH |  | Bark lignin conc. × DBH |  | Phloem lignin conc. × Bark lignin conc. |  |
| --- | --- | --- | --- | --- | --- | --- | --- | --- | --- | --- |
|  | Est. (95% CI) | p | Est. (95% CI) | p | Est. (95% CI) | p | Est. (95% CI) | p | Est. (95% CI) | p |
| *P. flexilis* | 0.14 (-0.02, 0.31) | 0.097 | -0.35 (-0.65, -0.03) | **0.037** | -0.03 (-0.10, 0.03) | 0.310 | 0.14 (-0.95, 0.74) | 0.739 | 0.02 (-0.01, 0.04) | 0.201 |
| *P. balfouriana* | 0.03 (-0.23, 0.31) | 0.812 | -0.32 (-2.38, 2.03) | 0.772 | -0.01 (-0.03, 0.02) | 0.493 | -0.24 (-1.06, 0.57) | 0.565 | -0.01 (-0.02, -0.00) | 0.055 |
| *P. longaeva* | -0.10 (-0.37, 0.15) | 0.448 | -0.11 (-1.18, 0.88) | 0.826 | 0.01 (-0.02, 0.05) | 0.403 | 0.15 (-0.84, 1.18) | 0.765 | -0.00 (-0.01, 0.01) | 0.892 |

## Discussion

Contrary to our expectations, *P. flexilis* exhibited the highest levels of constitutive phloem lignin relative to co-occurring *P. longaeva* and *P. balfouriana*, although there were no differences among the species in outer bark lignin. We also found no consistent relationship between phloem and outer bark lignin concentrations at the tree level. Because *P. flexilis* is considered more susceptible to MPB and produces greater numbers of offspring than *P. longaeva* and *P. balfouriana*, our results suggest that in these species constitutive lignin may not function as a direct defense against MPB attack or brood production. Our findings are similar to previous studies that showed phloem lignification did not differ among ash species (*Fraxinus spp.*) with varying resistance to the emerald ash borer (*Agrilus planipennis* Fair.) [74,75]. Although constitutive phloem lignin, as measured in our study, may not provide a significant defense, methyl jasmonate-induced lignification of *F. americana* and *F. pennsylvanica* phloem/outer bark was associated with resistance to the emerald ash borer [76]. The potential for induced lignification to act as an active defense in the *Pinus* species we sampled has not been investigated and should be part of future studies.

*Pinus flexilis* has consistently been found to have less constitutive and induced LMW specialized metabolites than other species, including *P. longaeva* and *P. balfouriana* at the sites sampled for this study [10], *P. contorta* and *P. ponderosa* [77], and the closely related bristlecone species *P. aristata* (Soderberg et al. in review). Although interspecific differences in selective pressure may have led to differences in investment in phloem specialized metabolite defenses [10, 78,79], our findings suggest an inverse relationship between phloem chemical defenses and lignification. In pines, phloem lignification is likely not selected in tandem with terpenoids that are known to provide defense against bark beetles [26,31], and our results suggest an investment tradeoff. In our study, *P. flexilis* had thinner phloem, but greater lignin concentrations and absolute abundance than *P. longaeva* and *P. balfouriana,* the latter two having thicker phloem. Moreover, *P. flexilis* with the thickest phloem had the lowest lignin concentrations, further suggesting a negative relationship between phloem thickness and lignification. The fact that outer bark lignin concentrations did not differ among the tree species but phloem concentrations did suggest that lignification within the phloem may be under different selective pressures relative to outer bark. Trait associations and underlying mechanisms facilitating phloem lignification may be unique to the functions of nutrient transport or defense against invading bacteria or pathogens, such as *Cronartium ribicola* J.C. Fisch., the causal agent of white pine blister rust, a serious disease of high elevation five needle pines [80]. However, this is highly speculative because the relative susceptibility to white pine blister rust among the three species in our study has not been investigated.

In summary, if defense against bark beetle attack were a strong selective driver for higher lignification in *Pinus*, higher lignin levels would be expected within both outer bark and phloem tissues of species considered less susceptible to MPB. This expectation is supported by prior work in *Picea* spp. [32,53,55,81]. However, our study demonstrated the opposite, i.e., that the more frequently attacked *P. flexilis* had greater phloem lignin levels relative to the less MPB-susceptible *P. longaeva* and *P. balfouriana*. Moreover, the species with the greatest constitutive phloem lignin concentrations, *P. flexilis*, was previously found to accumulate lower levels of constitutive LMW specialized metabolites than the other two species. While increased tissue lignification may have an additive effect in host defenses against MPB, there may be metabolic tradeoffs that are not accounted for between LMW specialized metabolites and lignin. Greater lignification within feeding tissues does not therefore appear to be generally adaptive as a defense against MPB. Interspecific differences in phloem but not outer bark lignin concentrations highlight that the benefits and costs of lignification are likely specific to phloem tissue.

## Acknowledgements

We thank Matt Hansen and Jim Vandygriff for providing assistance with field sample collection and Karen Mock for assistance in manuscript preparation.

## Supporting information

**S1 Fig. Lignin extracts of phloem and outer bark samples. All phloem samples were clear and colorless and therefore assumed pure (left vial). Outer bark samples were assumed to be pure when clear and colorless to light pink (right vial), but incompletely digested and/or contaminated when dark red (middle vial).**

**S1 Table. Model estimates testing for differences in phloem and bark lignin concentrations (g/mg FW) among sample sites of *P. flexilis, P. longaeva*, and *P. balfouriana* (see Table 1, Fig. 1). Effect size (Est.) and 95% confidence interval (95% CI) estimates between comparison samples are shown. P-values (p) presented describe the likelihood of statistical difference with values < 0.05 presented in bold.**

